# Kaempferol exerts anti-inflammatory effects by accelerating Treg development via AhR-mediated and PU.1/IRF4-dependent transactivation of the *Aldh1a2/*RALDH2 gene in dendritic cells

**DOI:** 10.1101/2024.02.21.581328

**Authors:** Miki Takahashi, Kazuki Nagata, Yumi Watanuki, Masaki Yamaguchi, Natsuki Minamikawa, Mayuka Katagiri, Weiting Zhao, Naoto Ito, Takuya Yashiro, Chiharu Nishiyama

## Abstract

Retinaldehyde dehydrogenase 2 encoded in the *Aldh1a2* gene (RALDH2) is expressed in intestinal dendritic cells (DCs) in mesenteric lymph nodes (MLNs) and plays a crucial role in Treg development by producing retinoic acid. In the present study, we screened food ingredients and identified kaempferol, a flavonoid, as the compound that most effectively up-regulated RALDH2 expression in DCs. The development of Foxp3^+^ cells from OT-II-derived naïve CD4^+^ T cells was enhanced by a co-culture with OVA peptide-pulsed bone marrow-derived DCs (BMDCs) in the presence of kaempferol or by a pre-treatment of BMDCs with kaempferol. The involvement of the aryl hydrocarbon receptor (Ahr) in the effects of kaempferol on *Aldh1a2* expression was demonstrated by *in vitro* experiments using siRNA and various agonistic or antagonistic compounds of AhR. AhR suppressed *Aldh1a2* gene expression and kaempferol inhibited the AhR activity by functioning as an antagonist of AhR. The kaempferol treatment also increased the protein levels of PU.1 and IRF4, which are essential for transcription of the *Aldh1a2* gene in DCs, and accelerated the subsequent recruitment of PU.1 to the *Aldh1a2* gene. The kaempferol-induced increase in PU.1 protein levels occurred in an AhR-dependent and transcription-independent manner, whereas the kaempferol treatment transactivated the *Irf4* gene in DCs. The frequency of DCs exhibiting RALDH2 activity in the MLNs of mice was increased by the intraperitoneally (i.p.) administration of an AhR antagonist or by the oral administration of astragalin, a 3-*O*-glucoside of kaempferol. We also observed an increase in Tregs in the Peyer’s patches of C57BL/6 mice i.p. administered kaempferol-treated BMDCs. We utilized an OVA-induced food allergy model of Balb/c mice to examine the effects of kaempferol *in vivo* and confirmed that the rapid decrease in body temperature and allergic diarrhea observed just after the OVA challenge were significantly suppressed in mice administered kaempferol.

## Introduction

Intestinal dendritic cells (DCs) in mesenteric lymph nodes (MLNs) specifically express the retinaldehyde dehydrogenase, RALDH2 (gene symbol *Aldh1a2*). RALDH2 converts retinal towards retinoic acid (RA), which functions as a ligand of the nuclear receptor RAR. DC-derived RA accelerates the development of Tregs by promoting the RAR-dependent transactivation of the *Foxp3* gene, which encodes a master transcription factor of Tregs. The essential role of RALDH2 in neonatal skin homeostasis was demonstrated in a recent study, which showed that the DC-specific deficiency of RALDH2 limited commensal-specific Treg generation in mice (1). Therefore, the up-regulation of gene expression and/or function of RALDH2 in DCs contribute to anti-inflammatory immunoresponses systemically.

In the present study, we performed screening for food ingredients that increase *Aldh1a2* mRNA levels in DCs, and the flavonoid, kaempferol was identified. Kaempferol and other flavonoids including quercetin, luteolin, and nobiletin, are the most vigorously studied polyphenols due to their beneficial effects in the host body, such as anticancer and anti-obesity activities. Previous studies, including ours, reported the anti-allergic effects of flavonoids targeting mast cells (2–8). Quercetin and luteolin, but not kaempferol, were recently identified as activators of the *Aldh1a2* promoter in an assay using the human monocyte cell line THP-1 carrying a reporter plasmid (9). Although a treatment with quercetin or luteolin increased *FOXP3* mRNA levels in human peripheral blood mononuclear cells (9), the molecular mechanisms by which flavonoids up-regulate *Aldh1a2* gene expression in monocyte lineages remain unclear.

In the present study, we revealed the involvement of the ligand-dependent transcription factor aryl hydrocarbon receptor (Ahr) and the hematopoietic cell-specific transcription factors PU.1 and IRF4. AhR is a ubiquitously expressed environmental sensor (particularly in barrier organs, including the skin and the mucosal tissues) and induces the expression of genes encoding drug-metabolizing enzymes, such as cytochrome C, in response to a ligand stimulation. So-called endocrine disruptors, including 2,3,7,8-tetrachlorodibenzo-p-dioxin (TCDD) and 3-methyl-cholanthrene, are well-known exogenous agonists of AhR. AhR also recognizes intestinal bacteria metabolites, such as indole-3-aldehyde, indole-3-acetic acid, indoxyl3-sulfate, and the tryptophan derivative indole-3-carbinol. PU.1 is an essential transcription factor for the development of DCs (10), and is required for the expression of DC-characteristic genes (11–13). We previously reported that PU.1 and its partner molecule IRF4 transactivated the *Aldh1a2* gene in DCs by binding to the *cis*-element on the gene (14). In the present study, we investigated the molecular mechanisms by which kaempferol up-regulates *Aldh1a2* transcripts in DCs, in which it regulates the function and/or expression of AhR, PU.1, and IRF4. We also examined the effects of kaempferol on the expression and/or function of RALDH2 in DCs and subsequent Treg development *in vitro* and *in vivo*.

## Materials and Methods

### Mice

C57BL/6 and Balb/c mice were purchased from Japan SLC (Hamamatsu, Japan). OT-II mice were from The Jackson Laboratory (USA). Mice were housed in a specific pathogen-free facility, and all animal experiments were performed in accordance with the guidelines of the Institutional Review Board of Tokyo University of Science. The present study was approved by the Animal Care and Use Committees of Tokyo University of Science: K22005, K21004, K20005, K19006, K18006, K17009, and K17012.

### Cells

Bone marrow-derived DCs (BMDCs) were generated from the bone marrow of C57BL/6 or Balb/c mice as previously described (12). BMDCs generated from C57BL/6 mice were pulsed with 25 μg/mL ovalbumin (OVA) peptide 323-339 (POV-3636-PI, Peptide Institute Inc., Osaka, Japan) for 1h. Cells were then washed and cultured with naïve CD4^+^ T cells isolated from the spleens of OT-II mice to induce the antigen-presenting cell-dependent activation of naïve CD4^+^ T cells. The MojoSort Mouse Naïve CD4^+^ T cell Isolation Kit (#480040, BioLegend, San Diego, CA, USA) was used for the isolation of naïve CD4^+^ T cells as previously described (15). MLN cells were collected from Balb/c mice and maintained in RPMI 1640 medium (#R8758, Sigma-Aldrich, St. Louis, USA) supplemented with 10% FBS, 100 U/mL penicillin, 100 μg/mL streptomycin, 100 μM 2-ME, 10 μM minimum non-essential amino acid solution, 1 mM sodium pyruvate, and 10 mM HEPES.

### Introduction of small interfering RNA (siRNA)

siRNAs for mouse AhR (#MSS201851), IDO1 (#MSS209789), IDO2 (#MSS205346), PU.1 (#MSS247676), and IRF4 (#MSS205501), and control siRNA (#12935-300, 12935-200) were purchased from Invitrogen (Carlsbad, CA, USA) and introduced into BMDCs by the electroporation system, Nucleofector 2b (Lonza, BASEL, Switzerland), using the Amaxa Mouse Dendritic Cell Nucleofector Kit (#VPA-1011, Lonza) as previously described (12, 16).

### Reagents

Kaempferol (#K0018, Tokyo Chemical Industry, Tokyo, Japan) and kaempferol-3-*O*-glucoside (Astragalin) (#1243S, SSX, Genay, Lyon area, France), TCDD (#D-404N, AccuStandard, Inc., New Haven, CT, USA), CH223191 (#16154, Cayman Chemical, Ann Arbor, MI, USA), IAId (#356-18121, Fujifilm Wako Chemicals, Osaka, Japan), L-kynurenine (kynurenine) (#K0016, Tokyo Chemical Industry), sesame oil (non-roasted) (#192-15382, Fujifilm Wako Chemicals), methyl cellulose 400 solution (#133-17815, Wako) were purchased from indicated source.

### Administration of compounds

Mice was intraperitoneally (i.p.) administered a total of 1 mg/kg of CH223191 or vehicle (sesame oil) 24 h before the analysis of RALDH2 activity in MLNs. After oral administration of 100 mg/kg/day kaempferol as a solution of the mixture of PEG400 and Tween80 (ratio 4:1) for 7 days, mice were sacrificed for the measurement of RALDH2 activity in MLNs. For 4 days, 15 mg/kg/day of astragalin or vehicle (0.5 w/v% methyl cellulose 400 solution) was orally given for the measurement of RALDH2 activity in the MLN. Mice were i.p. administered with 15 mg/kg/day kaempferol or vehicle (0.5 w/v% methyl cellulose 400 solution) in the OVA-induced food allergy.

### Quantification of mRNA levels

Total RNA extracted from BMDCs using the ReliaPrep RNA Cell Miniprep System (#Z6012, Promega, Madison, WI, USA) was reverse-transcribed to cDNA with ReverTra Ace qPCR RT Master Mix (#FSQ-201, TOYOBO, Osaka, Japan). Quantitative PCR was performed using a Step-One real-time PCR system (Applied Biosystems, Waltham, MA, USA) with THUNDERBIRD SYBR qPCR Mix (#QPS-201, TOYOBO) and primers. To measure the mRNA levels of *Actb*, *Gapdh*, *Hprt*, *aldh1a2* (RALDH2), *Ahr*, *Ido1*, *Ido2*, *Spi1* (PU.1), and *Irf4*, the following primers were used: *Actb* forward; 5’-agatgacccagatcatgtttgaga-3’, reverse; 5’-cacagcctggatggctacgta-3’, *Gapdh* forward; 5’-acgtgccgcctggagaa-3’, reverse; 5’-gatgcctgcttcaccacctt-3’, *Hprt* forward; 5’-tccattcctatgactgtagattttatcag-3’, reverse; 5’-aacttttatgtcccccgttgact-3’, *aldh1a2* forward; 5’-gacttgtagcagctgtcttcact-3’, reverse; 5’-tcacccatttctctcccatttcc-3’, *Ahr* forward; 5’-aatcccacatccgcatgatt-3’, reverse; 5’-tttgcaagaagccggaaaac-3’, *Ido1* forward; 5’-ctggcaaactggaagaaaaagg-3’, reverse; 5’-agaatgtccatgttctcgtatgtca-3’, *Ido2* forward; 5’-accaaaaggaacccagaagga-3’, reverse; 5’-ccggaaatgagatgatggtttc-3’, *Spi1* forward; 5’-atgttacaggcgtgcaaaatgg-3’, reverse; 5’-tgatcgctatggctttctcca-3’, *Irf4* forward; 5’-ccccattgagccaagcataa-3’, reverse; 5’-gcagccggcagtctgaga-3’.

### Flow cytometry

BMDCs and MLN cells were stained with the following Abs to identify DCs: anti-CD11c-PE/Cyanine 7 (clone; N418, TONBO Biosciences) and anti-I-A/I-E-PerCP (clone; M5/114.15.2, BioLegend). To detect Tregs in the co-cultured cells and Peyer’s patches, cell surface staining with anti-CD3ε-PerCP (clone; 145-2C11, BioLegend) and anti-CD4-FITC (clone; GK1.5, BioLegend) followed by intranuclear staining with an anti-Foxp3-APC Ab (clone; 3G3, TONBO Bioscience) using the Foxp3/Transcription Factor Staining Buffer Kit (#TNB-0607-KIT, TONBO Biosciences) was performed. CFSE (#65-0850-84, eBioscience Inc., San Diego, CA, USA) was used to analyze the proliferation of naïve CD4^+^ T cells. Fluorescence was detected by the MACS Quant Analyzer (Miltenyi Biotec) or FACS Lyric (Becton Dickinson) and analyzed with Flowjo software (Tomy Digital Biology, Tokyo, Japan).

### Measurement of enzymatic activity of RALDH2 by ALDEFLUOR assay

The enzymatic activity of RALDH2 in BMDCs and MLN cells was assesses using the ALDEFLUOR Kit (#01700, STEMCELL Technologies, Vancouver, BC, Canada). DCs were pre-gated as CD11c^+^/MHC II^high^ cells and RALDH2 activity was detected by a flow cytometric analysis.

### Chromatin immunoprecipitation (ChIP) assay

ChIP assays were performed as previously described (11, 14). Anti-PU.1 Ab (#sc-5949 X) was purchased from Santa Cruz Biotechnology (Santa Cruz, CA, USA). Goat IgG (#02-6202, Invitrogen) was used as the control Ab. The amount of chromosomal DNA was assessed by quantitative PCR using the StepOne Real-Time PCR system with the following primer sets: mouse RALDH2 promoter −1911/−1854 (forward primer, 5′-CCGGCCTCCCACTTAGGT-3′; reverse primer, 5′-AGCTCTCAAAAGTGGAGCAAA-3′).

### Western blotting

BMDCs (5.0 x 10^6^ cells) were collected after siRNA transfection and/or the kaempferol treatment. Subsequent steps, including the electrophoresis of cell lysates and transfer of proteins from acrylamide gels to membranes, were performed as previously described (11) using anti-PU.1 Ab (#sc-5949, Santa Cruz Biotechnology), anti-IRF4 goat polyclonal Ab (#sc-6059 X, Santa Cruz Biotechnology), and anti-β-actin murine mAb (#AC-15, Sigma-Aldrich).

### Induction and Assessment of OVA-induced food allergy

Female Balb/c mice (six-week old) were i.p. administered a mixture of 50 μg albumin from chicken egg white (OVA) (#A5503, Sigma-Aldrich) and 50 μL Alum (#77161, Thermo Fisher Scientific, Waltham, MA, USA) twice every other week. Two weeks after the second sensitization, mice were orally administered 50 mg OVA five times every other day. The rectal temperature of each mouse was measured every 10 minutes for 1 h just after the oral administration of OVA. The severity of diarrhea was assessed based on Table 2 and the number of stools was counted during the measurement of rectal temperatures. Symptoms were evaluated on the final day of the challenge.

### Statistical analysis

A two-tailed Student’s *t*-test was used to compare two samples, and a one-way ANOVA followed by Tukey’s multiple comparison test and Dunnett’s multiple comparison test were performed to compare more than three samples. *P* values < 0.05 were considered to be significant.

## Results

### Kaempferol up-regulates the gene expression and function of RALDH2 in DCs

We examined *Aldh1a2* mRNA levels in BMDCs treated with 39 food ingredients (Table 1) and identified kaempferol as the compound that most effectively increased *Aldh1a2* mRNA levels in DCs (Fig. 1A, Table 1). This effect of kaempferol was dependent on the dose applied (Fig. 1B) and was sustained for at least 48 h (Fig. 1C). A flow cytometry analysis revealed that increases in *Aldh1a2* mRNA levels were associated with enhancements in the enzyme activity of RALDH2 in BMDCs (Fig. 1D). We also confirmed that CD11c^+^/MHC II^high^ migDCs isolated from MLNs, which are a major population of RALDH2-expressing DCs (17), also exhibited higher RALDH2 activity in response to the treatment with kaempferol (Fig. 1E).

**Figure 1.**
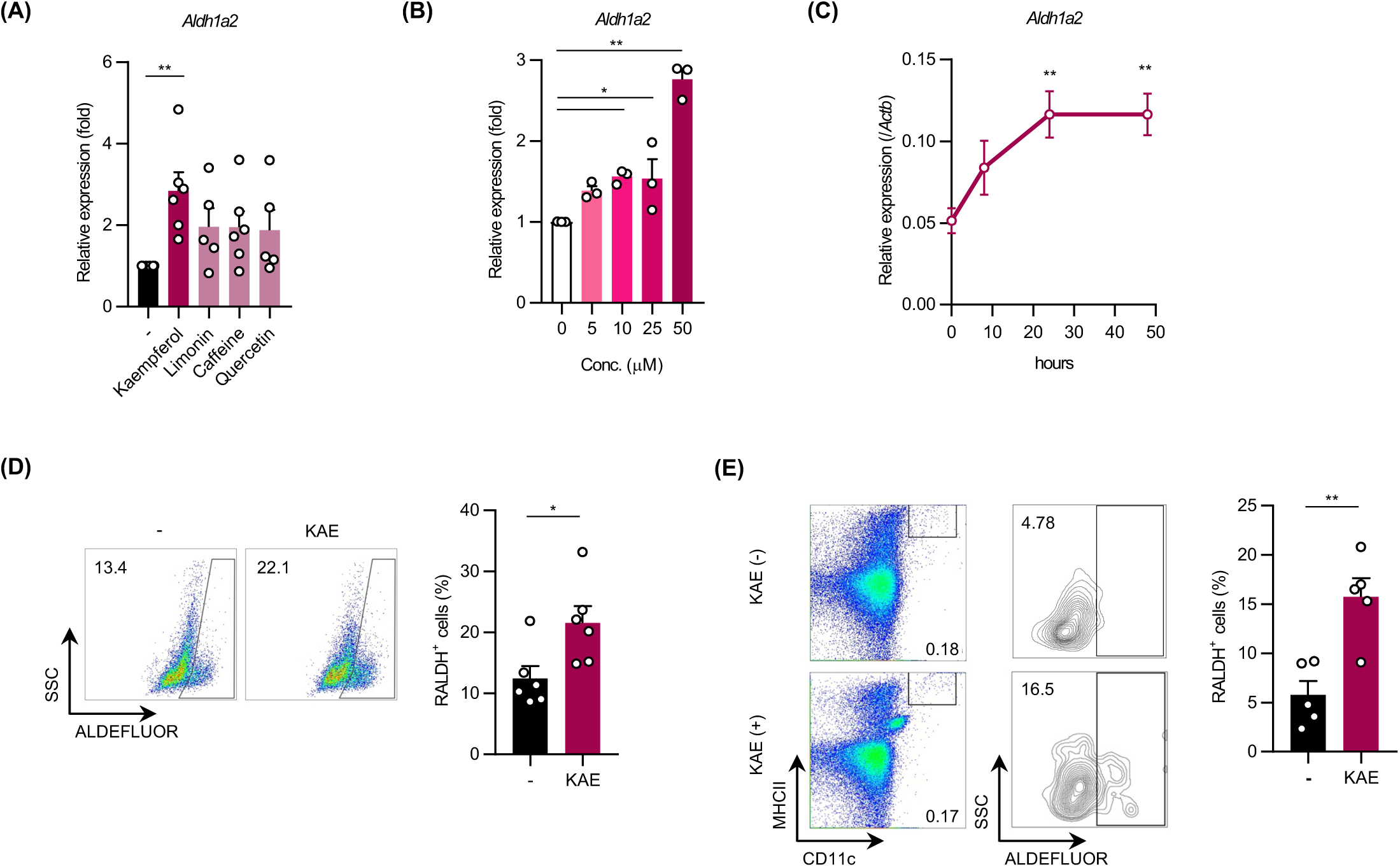
Kaempferol up-regulates the gene expression and function of RALDH2 in DCs. (**A**) *Aldh1a2* mRNA levels in food ingredient-treated BMDCs. BMDCs were incubated with the indicated food ingredients (50 μM) for 48h and mRNA levels were then assessed by qPCR. The top four candidates are shown in the graph and the remainder in Table 1. (**B**, **C**) Effects of kaempferol on the mRNA expression of *Aldh1a2*. BMDCs were incubated in the presence of kaempferol (5, 10, 25, and 50 μM) for 48 h (**B**) or 50 μM kaempferol for the indicated times (**C**). (**D**, **E**) The enzyme activity of RALDH2 in kaempferol-treated BMDCs (**D**) and CD11c^+^/MHC II^high^ migDCs from MLNs (**E**). BMDCs and MLN cells were treated with 50 μM kaempferol for 48 and 24 h, respectively. Data represent the mean ± SEM. Dunnett’s multiple comparison test (**A**-**C**) and two-tailed paired *t*-test (**D**, **E**) were used for statistical analyses. *, p < 0.05; n.s., not significant. Abbreviation: KAE, Kaempferol.

**Table 1.**
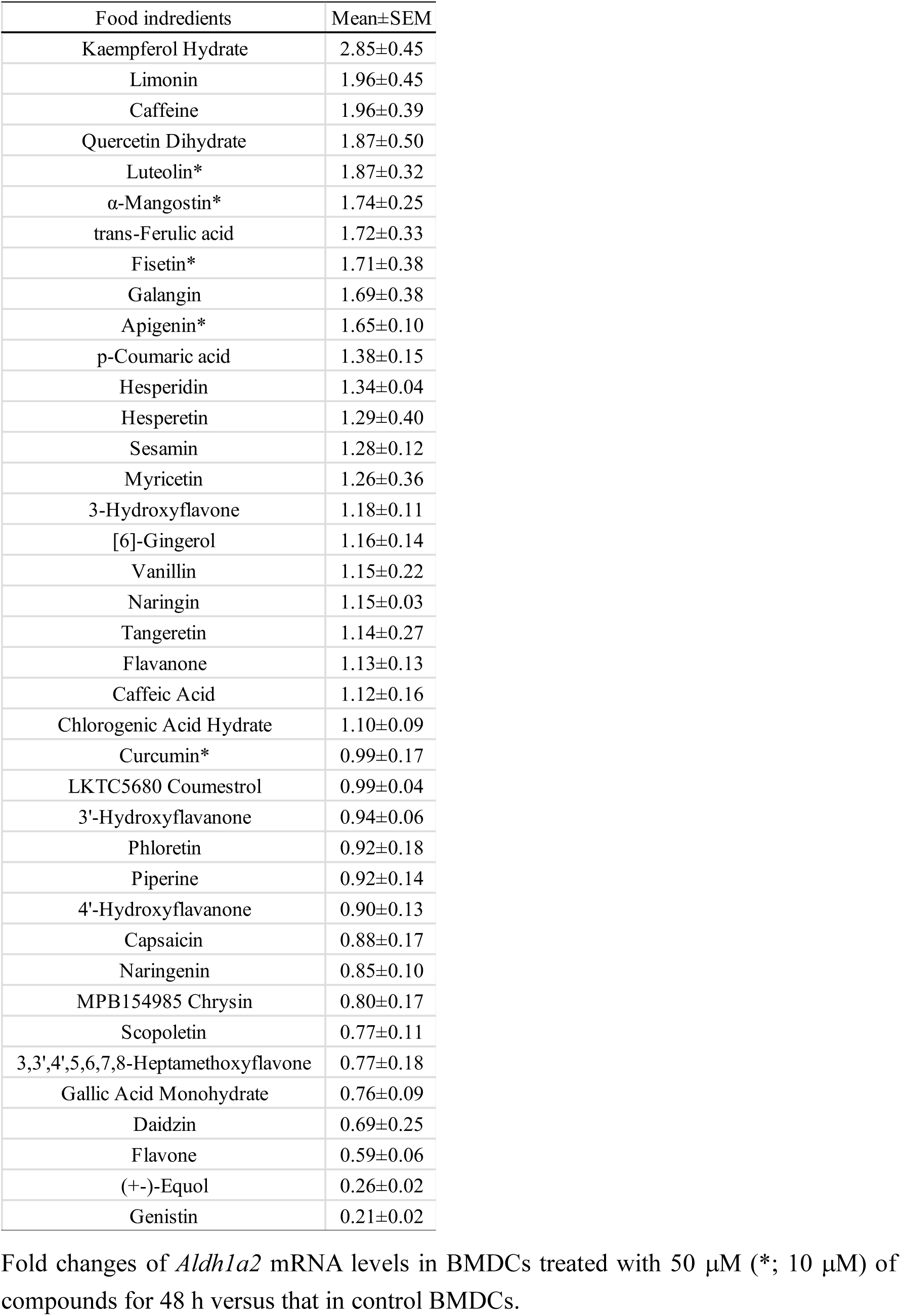
List of food ingredients subjected to screening.

**Table 2.** Criteria of diarrhea score.

| Score | Diarrhea |
| --- | --- |
| 0 | Hard |
| 1 | Soft |
| 2 | Loose |
| 3 | Muddy |
| 4 | Watery |

### Kaempferol-treated DCs accelerate Treg development

To establish whether the treatment with kaempferol affected the T cell development activity of DCs, we co-cultured peptide-pulsed BMDCs with naïve CD4^+^ T cells isolated from the spleens of OT-II mice. Under the condition that CD4^+^ T cells were divided by the co-culture with DCs, kaempferol significantly suppressed the division of T cells (Fig. 2A) and increased the frequency of Foxp3^+^ cells in CD4^+^ T cells (Fig. 2B). The increase in Foxp3^+^ cells was also observed in naïve CD4^+^ T cells, which were co-cultured with kaempferol-pretreated BMDCs (Fig. 2C), suggesting that kaempferol promoted Treg development by modulating the function of DCs.

**Figure 2.**
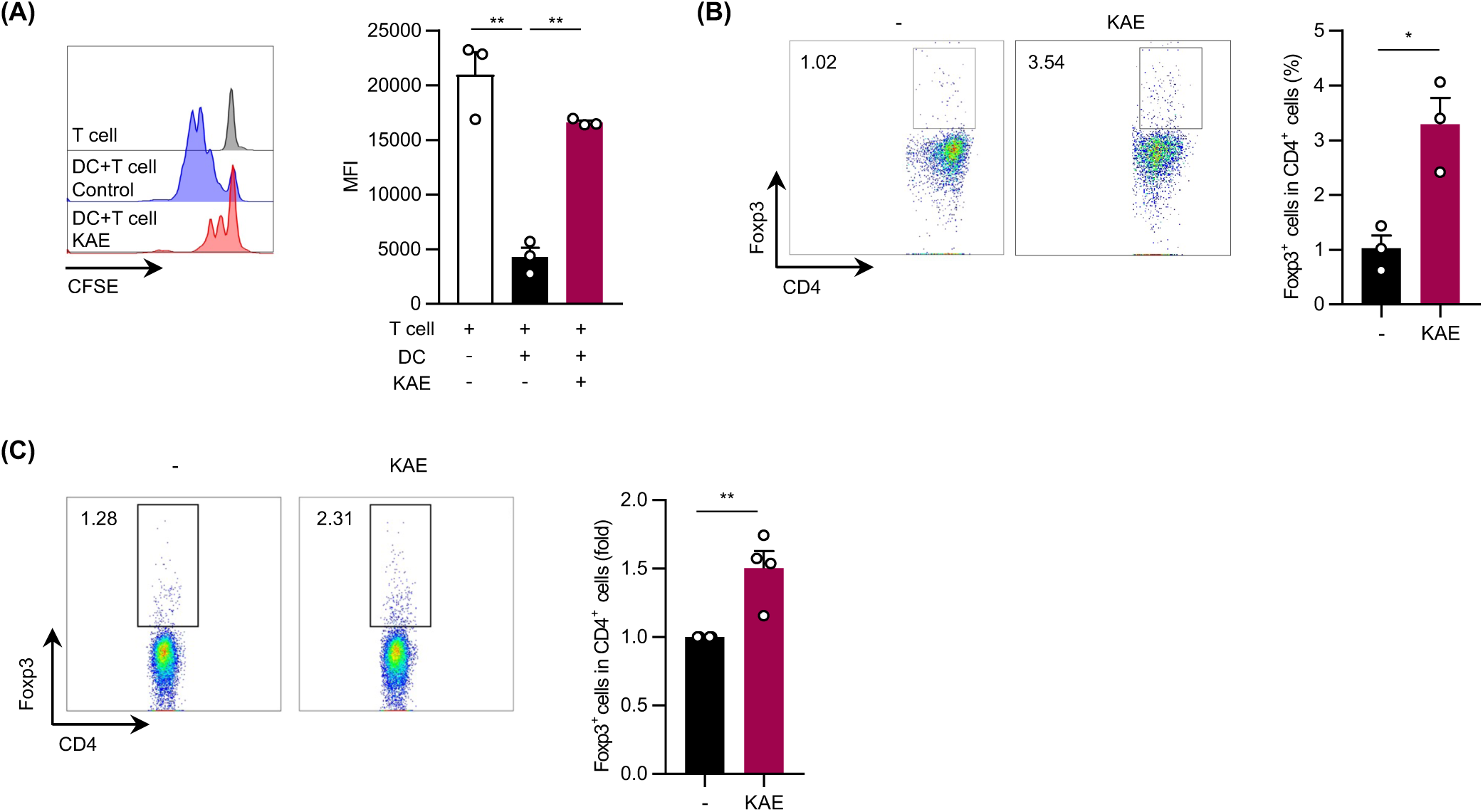
The kaempferol-induced up-regulation of RALDH2 in DCs accelerates Treg development. (**A**, **B**) The proliferation of naïve CD4^+^ T cells (**A**) and the frequency of Tregs (**B**). Naïve CD4^+^ T cells were co-cultured with OVA peptide-pulsed BMDCs in the presence of 50 μM kaempferol for 72 h (**A**, **B**). (**C**) The frequency of Tregs in CD4^+^ T cells co-cultured with kaempferol-pretreated BMDCs. BMDCs were cultured in the presence of 50 μM kaempferol for 48 h. Cells were washed and pulsed with the OVA peptide followed by a co-culture with naïve CD4^+^ T cells for 72 h. Data represent the mean ± SEM. Tukey’s multiple comparison test (**A**) and the two-tailed paired t-test (**B**, **C**) was used for statistical analyses. *, p < 0.05; n.s., not significant. Abbreviation: KAE, Kaempferol.

### Involvement of AhR in kaempferol-induced transactivation of the *Aldh1a2* gene

To investigate the molecular mechanisms by which kaempferol increased *Aldh1a2* mRNA levels in DCs, we performed knockdown experiments using siRNA targeting AhR, which has been reported to function as a receptor of kaempferol (18–20). As shown in Fig. 3A left, *Aldh1a2* mRNA levels were significantly higher in *Ahr* siRNA-transfected BMDCs than in control siRNA-transfected BMDCs, and the kaempferol-induced increase in *Aldh1a2* mRNA levels was canceled in *Ahr* siRNA-transfected BMDCs. The quantification of *Ahr* mRNA levels revealed the significant decrease of *Ahr* mRNA levels in *Ahr* siRNA-transfected BMDCs and a significant increase upon the kaempferol treatment (Fig. 3A right). The increase in *Aldh1a2* mRNA levels independent of the kaempferol treatment in *Ahr* siRNA-transfected BMDCs reflected the activity of RALDH2 (Fig. 3B). These results suggest the involvement of AhR in the kaempferol-mediated activation of gene expression and the function of RALDH2 in DCs as a negative regulator.

**Figure 3.**
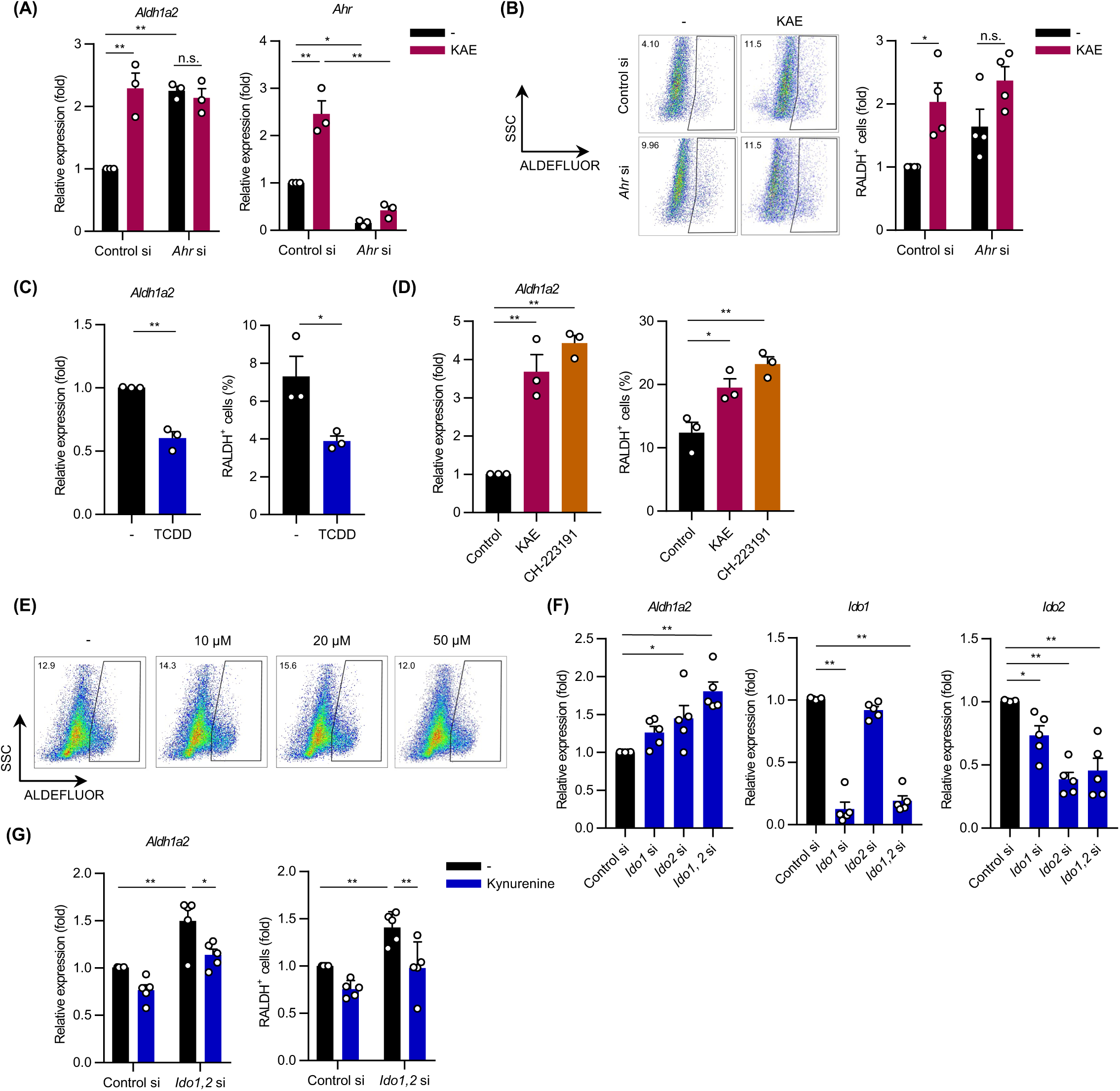
Involvement of AhR in the kaempferol-induced up-regulation of RALDH2. (**A**, **B**) The mRNA levels and activity of RALDH2 in *Ahr* siRNA-transfected BMDCs. *AhR* siRNA-transfected BMDCs were cultured in the presence of 50 μM kaempferol for 48 h. Cells were then harvested to assess the mRNA expression of *Aldh1a2* (**A** left) and *Ahr* (**A** right) and activity of RALDH2 (**B**). (**C**, **D**) *Aldh1a2* mRNA levels (left) and RALDH2 activity (right) in BMDCs treated with an agonist (**C**) and an antagonist (**D**) of AhR. BMDCs were cultured in the presence of 100 nM TCDD for 24 h (**C**) and 10 μM CH-223191 or 50 μM kaempferol for 48 h (**D**). (**E**) RALDH2 activity in BMDCs treated with kynurenine. BMDCs were treated with kynurenine (10, 20, and 50 μM) for 24 h and then subjected to the ALDEFLOUR assay. A typical result is shown, and similar result was obtained in another independent experiment. (**F**) *Aldh1a2* mRNA levels in BMDCs with the knockdown of IDO. BMDCs were transfected with *Ido1* and/or *Ido2* siRNA and harvested 48 h later. (**G**) The mRNA levels and activity of RALDH2 in BMDCs with the knockdown of *Ido*. BMDCs transfected with *Ido1* and *Ido2* siRNA were cultured in the presence of 50 μM kynurenine for 48 h. Data represent the mean ± SEM. Tukey’s multiple comparison test (**A**, **B**, **G**), the two-tailed paired *t*-test (**C**), and Dunnett’s multiple comparison test (**D**, **F**) were used for statistical analyses. *, p < 0.05; n.s., not significant. Abbreviation: KAE, Kaempferol.

To examine the roles of AhR in the kaempferol-induced transactivation of the *Aldh1a2* gene, we treated BMDCs with TCDD, an agonist of AhR (Fig. 3C), and CH-223191, an antagonist of AhR (Fig. 3D). As expected, the mRNA level and enzymatic activity of RALDH2 were reduced by the TCDD treatment (Fig. 3C), but were significantly enhanced by the treatment with CH-223191, similar to kaempferol (Fig. 3D).

The results in Fig. 3A showing an increase in *Aldh1a2* mRNA levels following the knockdown of *Ahr* indicated that the suppressive functions of AhR in the expression of the *Aldh1a2* gene were constitutively activated in cultured BMDCs, suggesting the presence of endogenous agonist(s). We considered kynurenine, an agonist of AhR (21, 22), as a candidate DC-derived endogenous agonist involved in the kaempferol-mediated regulation of the *Aldh1a2* gene. When various concentrations of kynurenine were added to the culture media of BMDCs, RALDH2 activity was not affected (Fig. 3E). We then performed the knockdown of *Ido1* and/or *Ido2*, which encode indole 2,3-dioxygenase 1 and 2, respectively, the rate-limiting enzymes of kynurenine synthesis from tryptophan in DCs. When the mRNA levels of *Ido1* and *Ido2* were significantly reduced by siRNA transfection, *Aldh1a2* mRNA levels increased in DCs (Fig. 3F). This increase was attenuated by the addition of kynurenine to the culture media of BMDCs (Fig. 3G). These results demonstrate that kaempferol up-regulated *Aldh1a2* gene expression by antagonizing AhR, which suppressed *Aldh1a2* gene expression with constitutive activation by endogenously synthesized kynurenine in DCs.

### Roles of PU.1 and IRF4 in kaempferol-induced transactivation of the *Aldh1a2* gene

We previously reported that PU.1 and IRF4 transactivated the *Aldh1a2* gene by binding to the enhancer element around −2kb of the *Aldh1a2* gene in DCs (14). To clarify the involvement of PU.1 and IRF4 in kaempferol-induced increases in *Aldh1a2* mRNA levels, we examined the mRNA and protein levels of these transcription factors in kaempferol-treated BMDCs by qPCR (Fig. 4A) and Western blotting (Fig. 4B), respectively. Although a significant difference was not observed in the mRNA level of *Spi1*, the levels of *Irf4* mRNA, the IRF4 protein, and PU.1 protein in DCs were significantly increased by the treatment with kaempferol (Fig. 4A, 4B). We also revealed that the recruitment of PU.1 to the enhancer region of the *Aldh1a2* gene was significantly enhanced in kaempferol-treated DCs (Fig. 4C). Furthermore, the kaempferol-induced increase in *Aldh1a2* mRNA levels was abolished in DCs following the knockdown of *Spi1* or *Irf4* by their siRNAs (Fig. 4D).

**Figure 4.**
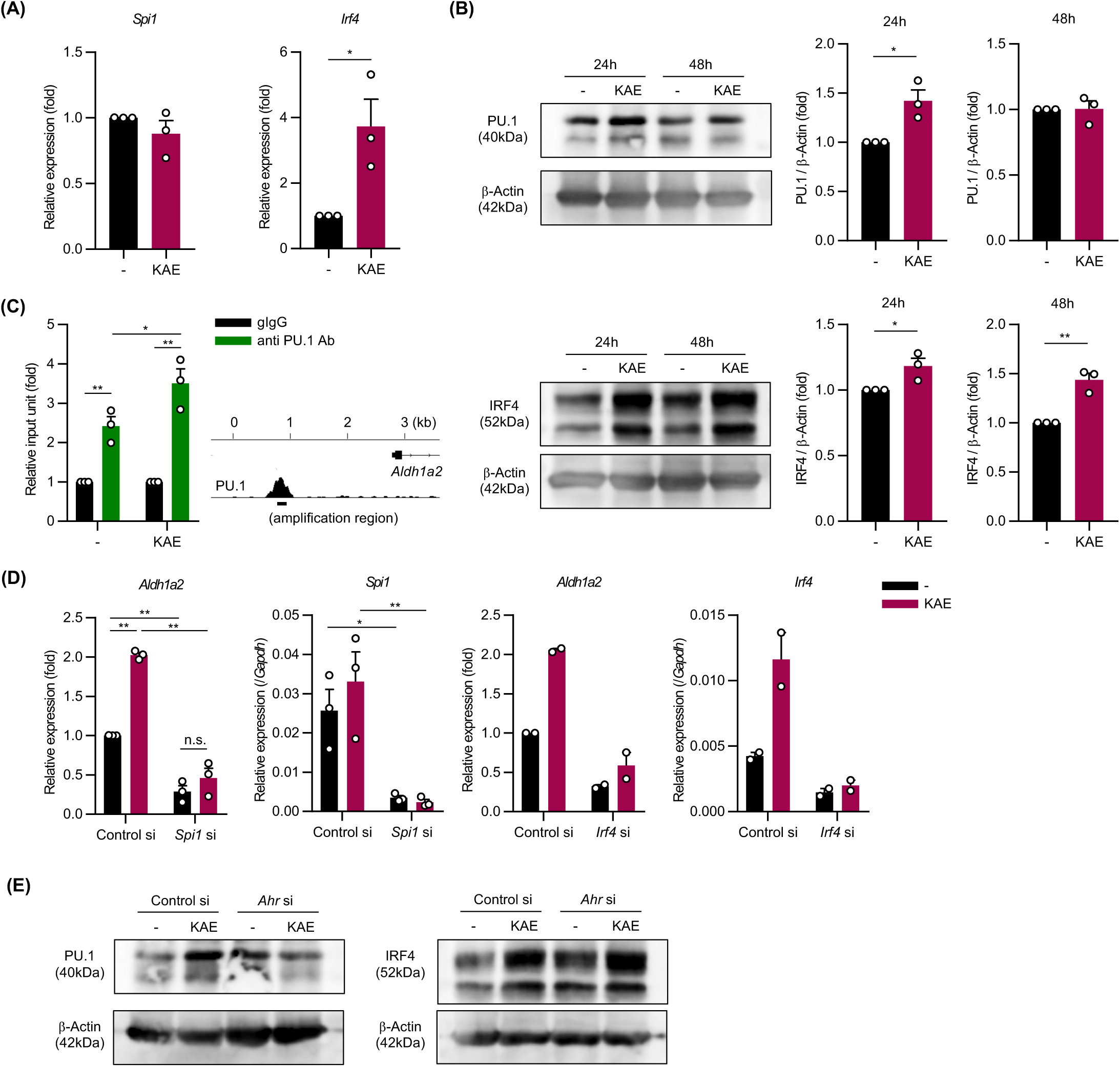
Effects of the kaempferol treatment on transcription factors PU.1 and IRF4. (**A**, **B**) The mRNA (**A**) and protein (**B**) levels of PU.1 and IRF4 in BMDCs. BMDCs were cultured in the presence of 50 μM kaempferol for 24 - 48 h. Data of three independent experiments were quantified by calculating the band intensity, and shown as ratio to that of control (kaempferol-non-treated) BMDCs. (**C**) Genomic DNA levels in BMDCs immunoprecipitated by ChIP assays using anti-PU.1 Ab were assessed by qPCR targeting the enhancer region of the *Aldh1a2* gene. BMDCs were incubated in the presence or absence of 50 μM kaempferol for approximately 24 h. (**D**) *Aldh1a2* mRNA levels in siRNA-transfected BMDCs. BMDCs transfected with *Spi1* or *Irf4* siRNA were cultured in the presence of 50 μM kaempferol for 48 h. (**E**) PU.1 and IRF4 protein levels in *Ahr* siRNA-transfected BMDCs. *Ahr* siRNA- or control siRNA-transfected BMDCs were cultured in the presence of kaempferol for 24 h. Data represent the mean ± SEM. The two-tailed paired *t*-test (**A**, **B**) and Tukey’s multiple comparison test (**C**, **D**) were used for statistical analyses. *, p < 0.05; n.s., not significant. Abbreviation: KAE, Kaempferol.

We also investigated PU.1 and IRF4 protein levels of in *Ahr* siRNA-transfected DCs to reveal the roles of AhR in kaempferol-induced increases in PU.1 and IRF4. Western blot profiles showed that kaempferol-induced increases in PU.1 protein levels observed in control siRNA-transfected DCs were eliminated by the knockdown of *Ahr*, whereas kaempferol-induced increases in IRF4 protein levels were not, and *Ahr* knockdown tended increase IRF4 protein levels (Fig. 4E).

These results indicate that the kaempferol treatment increased the protein levels of IRF4 and PU.1, which play crucial roles in the transcription of the *Aldh1a2* gene, in DCs via distinct mechanisms; namely, by transactivating the *Irf4* gene and by increasing the level of PU.1 in an AhR-dependent and transcription-independent manner.

### Effects of kaempferol on OVA-induced food allergy in mice

To examine the effects of AhR antagonization on RALDH2 activation in DCs *in vivo*, we analyzed RALDH2 activity in DCs isolated from the MLNs of mice, which were i.p. administered 1 mg/kg CH-223191 24 h before. As shown in Fig. 5A, the frequency of RALDH2-positive cells in migDCs was significantly increased by the administration of the AhR antagonist. To investigate whether kaempferol-treated DCs accelerated Treg development *in vivo*, we i.p. administered BMDCs pre-incubated with or without kaempferol. A flow cytometry analysis showed a significantly higher frequency of Fopx3^+^/CD4^+^ T cells in the Peyer’s patches from mice injected with kaempferol-treated BMDCs than from those injected with non-treated BMDCs (Fig. 5B).

**Figure 5.**
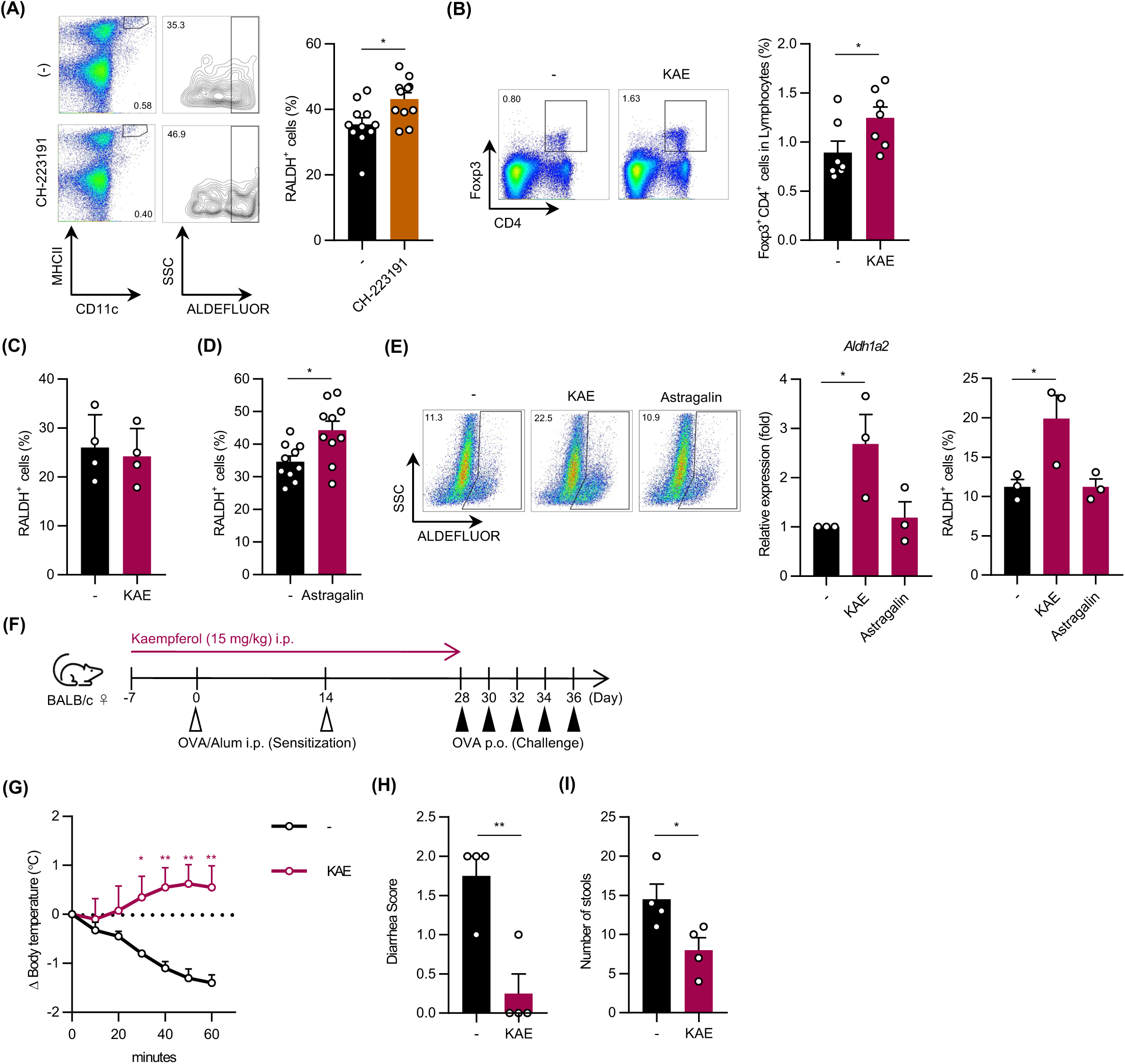
Effects of kaempferol on RALDH2-related immune regulation in vivo. (**A**) RALDH2 activity in CD11c^+^/MHC II^high^ migDCs isolated from MLNs. CH-223191 (1 mg/kg) was intraperitoneally (i.p.) administered 24 h before the analysis. (**B**) The frequency of CD4^+^/Fopx3^+^ cells in the Peyer’s patches isolated from BMDC-injected mice. BMDCs were pre-incubated with or without 50 μM kaempferol for 48 h and 0.3-2.0×10^6^ cells were i.p. administered to each mouse 4 days before the analysis. (**C**, **D**) RALDH2 activity in CD11c^+^/MHC II^high^ migDCs isolated from MLNs. Mice were orally administered 100 mg/kg kaempferol for 7 days (**C**) or 15 mg/kg astragalin for 4 days (**D**). *Aldh1a2* mRNA levels and RALDH2 activity in astragalin-treated BMDCs. BMDCs were incubated with 50 μM astragalin or kaempferol for 48 h. (**F**-**I**) The OVA-induced food allergy model. A scheme of the experiment (**F**). The decrease in body temperature (**G**), diarrhea scores (**H**), and the number of stools (**I**) after the OVA challenge (on Day 36) were evaluated. Data represent the mean ± SEM. The two-tailed paired *t*-test (**A**-**D**, **G**-**I**) and Dunnett’s multiple comparison test (**E**) were used for statistical analyses. *, p < 0.05; n.s., not significant. Abbreviation: KAE, Kaempferol.

We analyzed RALDH2 activity in migDCs isolated from the MLNs of mice orally administered 100 mg/kg/day kaempferol for 7 days to investigate the effects of kaempferol intake *in vivo*. The frequency of RALDH2^+^ cells in the DCs of mice, which were sacrificed on day 8 (24 h after the last administration), was similar between kaempferol-treated mice and their controls (Fig. 5C). Previous findings showed that the oral intake of flavonoid glycosides exerted stronger biological effects than that of flavonoid aglycones by maintaining a higher serum concentration (23); therefore, we used a glycoside instead of an aglycon in the oral administration experiment. The oral administration of 15 mg/kg/day astragalin for 4 days, which is the 3-*O*-glucoside of kaempferol, enhanced RALDH2 activity (Fig. 5D), but did not increase the gene expression or enzymatic activity of RALDH2 *in vitro* (Fig. 5E).

To investigate the effects of the kaempferol treatment on immune-related diseases, we utilized an OVA-induced food allergy model of Balb/c mice. The i.p. administration of 15 mg/kg/day kaempferol was initiated 7 days prior to the first sensitization with OVA and was continued until the day before the first challenge (Fig. 5F). Under the condition that control mice not treated with kaempferol showed allergic symptoms, including a decreased body temperature due to anaphylaxis, diarrhea, and increased frequency of defecation, the kaempferol treatment significantly suppressed OVA-induced anaphylaxis (Fig. 5G), diarrhea scores (Fig. 5H), and the number of stools (Fig. 5I).

## Discussion

In the present study, we identified kaempferol from the screening of food ingredients that up-regulate *Aldh1a2* gene expression in DCs. *In vitro* experiments indicated that kaempferol increased the mRNA levels and enzymatic activity of RALDH2 by antagonizing AhR in DCs. In addition, kaempferol increased the protein levels of IRF4 and PU.1 in a transcription-mediated manner and in AhR-dependent manner, respectively. We also found that Treg development activity was enhanced in kaempferol-treated DCs. Previous studies demonstrated the essential roles of GM-CSF in RALDH2 expression in mouse and human DCs (24, 25). GM-CSF stimulation markedly increased *Aldh1a2* mRNA levels and RALDH2 enzymatic activity in mouse splenic DCs (24). *Aldh1a2* mRNA levels were higher in GM-CSF-induced BMDCs than in Flt3L-induced BMDCs (24). Regarding human DCs, Vitamin D3 (VD3) synergistically transactivates the *Aldh1a2* gene in GM-CSF-maintained human CD1c^+^ DCs (25). Furthermore, GM-CSF increased the mRNA and protein levels of IRF4 as well as PU.1 protein binding to the *Aldh1a2* gene in Flt3L-induced BMDCs (14). Increases in IRF4 transcription and PU.1 recruitment in DCs were also observed in kaempferol-treated DCs in the present study. Kaempferol may modulate GM-CSFR-mediated signal transduction in GM-CSF-induced BMDCs. Alternatively, the kaempferol treatment may up-regulate GM-CSF expression in BMDCs, which subsequently stimulate DCs in an autocrine manner. Although we used GM-CSF-induced BMDCs in the present study, an investigation using Flt3L-induced BMDCs and splenic DCs, which are more sensitive to a GM-CSF stimulation, is needed to examine the effects of kaempferol on RALDH2 expression in DCs. It is important to note the difference between the effects of kaempferol treatment and GM-CSF stimulation on PU.1 protein levels in DCs. PU.1 protein levels increased in kaempferol-treated BMDCs in the present study, whereas GM-CSF was previously shown to accelerate the recruitment of PU.1 to the *Aldh1a2* gene without affecting PU.1 protein levels in DCs (14). The increase in PU.1 protein levels noted in kaempferol-treated DCs was dependent on AhR. Although AhR is a ubiquitous protein, its expression is higher in barrier organs, including the skin and the mucosal tissues. Among human myeloid lineages, the AhR protein is expressed at high levels in Langerhans cells (LCs) and at intermediate levels in monocytes, but at low to undetectable levels in granulocytes and expanded progenitors (26). The AhR ligand, VAF347, which suppresses allergic inflammation and allograft rejection, inhibited LC development by down-regulating PU.1 expression (26). Although VAF347 decreased *SPI1* mRNA levels and kaempferol increased PU.1 protein levels without affecting *Spi1* mRNA levels, these findings suggest the involvement of AhR in the development and function of the DC lineage through the regulation of PU.1. Since PU.1 transactivates several genes characteristic of DCs, including MHC II (11), CD80 and CD86 (12), and PD-L2 (27), kaempferol is expected to regulate DC-dependent immune responses by modulating the expression of various genes in DCs. Furthermore, AhR has been shown to regulate commitment between Th17 and Tregs (28). Although kaempferol did not affect Treg development from naïve CD4^+^ T cells following a stimulation with anti-CD3 and anti-CD28 Abs, namely DC-independent conditions, in our preliminary experiment (data not shown), the effects of kaempferol on other immune-related cells warrant further study to clarify its potency as an immunomodulator.

In addition to kaempferol, apigenin, luteolin, fisetin, galangin, and quercetin, all of which are flavonoids, exhibited high transactivation activity for the *Alldh1a2* gene in our screening (Fig. 1 and Table I). Based on the structures of flavonoids, they may prefer to bind to the ligand-binding pocket of AhR as antagonists. In addition, we hypothesized that the antioxidant activities of flavonoids contributed to their anti-inflammatory effects. Previous studies indicated that kaempferol activated NRF2 (29, 30), which is a master transcription factor of antioxidant stress and its pathway is often identified as a target of phytochemicals exerting anti-inflammatory effects (31, 32). Our preliminary experiment using *Nrf2*^-/-^ BMDCs showed that enhanced Treg development in a coculture with kaempferol-treated DCs was reduced by a NRF2 deficiency, whereas kaempferol-induced increases in *Aldh1a2* mRNA levels were not decreased in *Nrf2*^-/-^ DCs (data not shown), suggesting the NRF2 is not involved in the up-regulation of RALDH2 in kaempferol-treated DCs.

The roles of RA in RALDH2 expression in DCs have been investigated (24, 33). RA increased *Aldh1a2* mRNA levels in Flt3L-induced BMDCs, and an RAR antagonist partially inhibited GM-CSF-induced increases in *Aldh1a2* mRNA levels in Flt3L-induced DCs and splenic DCs (24). The presence of RA during the development of DCs up-regulated *Aldh1a2* expression and this was accompanied by the enforced expression of CCR9 and TGF-β (33). As described above, VD3 up-regulated RALDH2 expression in human DCs (25). The molecular mechanisms by which these fat-soluble vitamins affect RALDH2 expression in DCs remain largely unknown. Further studies on the relationships between RAR, VDR, AhR, and PU.1/IRF4 in DCs are needed to clarify the molecular mechanisms underlying the immuno-modulatory effects of food ingredients.

## Authorship Contribution

M.T. performed experiments, analyzed data, and prepared figures; K.N, analyzed data, prepared figures, and wrote the manuscript, Y.W., and M.Y. performed experiments, and analyzed data; N.M., M.K., W.Z., N.I., and T.Y. performed experiments; C.N. supervised, designed research, and wrote the manuscript.

## Acknowledgments

We thank the members of the Laboratory of Molecular Biology and Immunology, Department of Biological Science and Technology, Tokyo University of Science for their constructive discussions and technical support. We greatly appreciate the consideration and support from Dr. Kimihiko Yasuda, Dr. Masako Yasuda, and the late Ms. Yayoi Yasuda.

## Abbreviations

AhR, aryl hydrocarbon receptor; BM, bone marrow-derived; ChIP, chromatin immunoprecipitation; DC, dendritic cell; IAId, indole-3-aldehyde; i.p., intraperitoneal; MLN, mesenteric lymph node; OVA, ovalbumin; RALDH, retinaldehyde dehydrogenase; RA; retinoic acid; siRNA, small interfering RNA; TCDD, 2,3,7,8-tetrachlorodibenzo-p-dioxin; VD3, vitamin D3.

